# Repurposing the Ebola and Marburg Virus Inhibitors Tilorone, Quinacrine and Pyronaridine: In vitro Activity Against SARS-CoV-2 and Potential Mechanisms

**DOI:** 10.1101/2020.12.01.407361

**Authors:** Ana C. Puhl, Ethan James Fritch, Thomas R. Lane, Longping V. Tse, Boyd L. Yount, Carol Queiroz Sacramento, Tatyana Almeida Tavella, Fabio Trindade Maranhão Costa, Stuart Weston, James Logue, Matthew Frieman, Lakshmanane Premkumar, Kenneth H. Pearce, Brett L. Hurst, Carolina Horta Andrade, James A. Levi, Nicole J. Johnson, Samantha C. Kisthardt, Frank Scholle, Thiago Moreno L. Souza, Nathaniel John Moorman, Ralph S. Baric, Peter Madrid, Sean Ekins

## Abstract

SARS-CoV-2 is a newly identified virus that has resulted in over 1.3 M deaths globally and over 59 M cases globally to date. Small molecule inhibitors that reverse disease severity have proven difficult to discover. One of the key approaches that has been widely applied in an effort to speed up the translation of drugs is drug repurposing. A few drugs have shown *in vitro* activity against Ebola virus and demonstrated activity against SARS-CoV-2 *in vivo*. Most notably the RNA polymerase targeting remdesivir demonstrated activity *in vitro* and efficacy in the early stage of the disease in humans. Testing other small molecule drugs that are active against Ebola virus would seem a reasonable strategy to evaluate their potential for SARS-CoV-2. We have previously repurposed pyronaridine, tilorone and quinacrine (from malaria, influenza, and antiprotozoal uses, respectively) as inhibitors of Ebola and Marburg virus *in vitro* in HeLa cells and of mouse adapted Ebola virus in mouse *in vivo*. We have now tested these three drugs in various cell lines (VeroE6, Vero76, Caco-2, Calu-3, A549-ACE2, HUH-7 and monocytes) infected with SARS-CoV-2 as well as other viruses (including MHV and HCoV 229E). The compilation of these results indicated considerable variability in antiviral activity observed across cell lines. We found that tilorone and pyronaridine inhibited the virus replication in A549-ACE2 cells with IC_50_ values of 180 nM and IC_50_ 198 nM, respectively. We have also tested them in a pseudovirus assay and used microscale thermophoresis to test the binding of these molecules to the spike protein. They bind to spike RBD protein with K_d_ values of 339 nM and 647 nM, respectively. Human C_max_ for pyronaridine and quinacrine is greater than the IC_50_ hence justifying *in vivo* evaluation. We also provide novel insights into their mechanism which is likely lysosomotropic.

## Introduction

At the time of writing (November 2020) we are in the midst of a global health crisis caused by a new virus that originated in Wuhan, China in late 2019 and which has already generated considerable economic and social hardship. A new coronavirus called severe acute respiratory coronavirus 2 (SARS-CoV-2) shares aspects of pathology and pathogenesis with closely related SARS-CoV (Coronaviridae Study Group of the International Committee on Taxonomy of, 2020; Wu et al., 2020) and Middle East Respiratory Syndrome coronavirus (MERS-CoV) (Liu et al., 2020b), which also belong to the same family of *Betacoronavirus*. These viruses cause highly pathogenic respiratory infection that may lead to considerable morbidity, mortality and the broad range of clinical manifestations associated with SARS-CoV-2 which has been collectively called 2019 coronavirus disease (COVID-19) (WHO). SARS-CoV-2 infection may result in cough, loss of smell and taste, respiratory distress, pneumonia and extrapulmonary events characterized by a sepsis-like disease that require hospitalization, and may lead to death (Pan et al., 2020). Similar with SARS-CoV, SARS-CoV-2 directly interacts with angiotensin converting enzyme 2 (ACE2) receptor in host cell types (Brann et al., 2020; Sungnak et al., 2020; Whitcroft and Hummel, 2020). Because COVID-19 is established as new public health problem and vaccines are unlikely to eradicate animal reservoirs of SARS-CoV-2, inhibition of key events during the viral life cycle could pave the way for repurposed drugs.

Indeed, SARS-CoV-2 spread rapidly worldwide prompting the World Health Organization to declare the outbreak a pandemic, with more than 1.5 million cases confirmed in less than 100 days. At the time of writing there are over 59 million confirmed cases (WHO, 2020). The high infection rate has caused considerable stress on global healthcare systems leading to over 1.3M deaths from COVID-19 and the USA has reported the largest number of fatalities (WHO, 2020). Epidemic and pandemic disease outbreaks have intensified in recent years and this will require various small molecule therapeutic interventions to be developed.

There have been many efforts globally to screen and identify drugs for SARS-CoV-2 and there are currently clinical trials using existing drugs that are being repurposed. One early success was the RNA polymerase inhibitor remdesivir, which had previously been in a clinical trial for Ebola virus (EBOV) (Mulangu et al., 2019), while also demonstrating inhibition of MERS activity in rhesus macaques (de Wit et al., 2020) and against many SARS-like coronaviruses, including SARS-CoV-2 in primary human cells and *in vivo* (Pruijssers et al., 2020; Sheahan et al., 2017). Remdesivir demonstrated activity in Vero cells (Pruijssers et al., 2020; Wang et al., 2020a), human epithelial cells and in Calu-3 cells (Pruijssers et al., 2020) infected with SARS-CoV-2, which justified further testing in the clinic. This drug was then the subject of numerous clinical trials globally (Beigel et al., 2020a; Lamb, 2020; Wang et al., 2020b). These included a randomized double-blind, placebo-controlled multicenter trial that demonstrated that remdesivir reduced the days to recovery (Wang et al., 2020b) although adverse events were also higher in treated versus placebo groups. Regardless, it quickly received an emergency use authorization (Lamb, 2020). Recent double-blind, randomized, placebo-controlled trial in adults hospitalized with COVID-19 and had evidence of lower respiratory tract infection, demonstrated remdesivir was superior to placebo in shortening the time to recovery (Beigel et al., 2020a). Most recently a large multicenter SOLIDARITY trial showed no efficacy for hospitalized patients with COVID-19, indicating that this drug may be more useful early in infection. This drug was also recently approved by the FDA in October 2020.

There have also been notable failures including hydroxychloroquine which was also initially identified as active in Vero cells *in vitro* (Liu et al., 2020a) but repeatedly failed spectacularly in numerous clinical trials (Boulware et al., 2020; Roomi et al., 2020).

Still, repurposing drugs represents possibly the fastest way to identify a drug and bring it to the clinic with fewer potential hurdles if it is already an approved drug or clinical candidate (Baker et al., 2018; Guy et al., 2020). There have been several large-scale high throughput screens, one used Huh-7 cells and tested 1425 compounds, identifying 11 with activity IC_50_ < 1 μM (Mirabelli et al., 2020). A screen of 1528 compounds lead to 19 hits in Vero cells including 4 with IC_50_’s of ~1 μM (Yuan et al., 2020). A screen of the Prestwick library in hPSC lung organoids identified 3 hits (Han et al., 2020). 12,000 clinical stage or FDA approved compounds in the ReFRAME library screened using Vero cells led to 21 hits (Riva et al., 2020). To date we have collated well over 500 drugs that have *in vitro* data from the various published *in vitro* studies against this virus (Jeon et al., 2020b; Jin et al., 2020; Liu et al., 2020a; Wang et al., 2020a) and used these to build a machine learning model that was used to select additional compounds for repurposing and testing (Gawriljuk et al., 2020). Several of these molecules had also previously demonstrated *in vitro* activity against the Ebola virus (EBOV). For example, a machine learning model was previously used to identify tilorone, quinacrine and pyronaridine tetraphosphate (Fig.1) (Ekins et al., 2015a) and all inhibited EBOV and Marburg *in vitro* and they showed significant efficacy in the mouse-adapted EBOV (ma-EBOV) model (Ekins et al., 2020; Ekins et al., 2018a; Lane et al., 2019a; Lane et al., 2019c). All of these molecules are also lysosomotropic (Lane and Ekins, 2020) which may contribute to their effect as possible entry inhibitors. Pyronaridine tetraphosphate is currently used as an antimalarial in combination with artesunate (Pyramax®) and has also shown significant activity in the guinea pig adapted model of EBOV infection (Lane et al., 2020c). Tilorone is structurally different and is used in eastern Europe as an antiviral for influenza (Ekins and Madrid, 2020c). Tilorone has also demonstrated *in vitro* inhibition of MERS (Ekins et al., 2020; Ekins and Madrid, 2020b), SARS-CoV-2 and has shown a low μM IC_50_ (Jeon et al., 2020a). Most recently tilorone, quinacrine and pyronaridine tetraphosphate were demonstrated to bind to the EBOV glycoprotein. The K_d_ values for pyronaridine (7.34 μM), tilorone (0.73 μM) and quinacrine (7.55 μM) possessed higher affinity than previously reported compounds that were found to bind to this protein (Lane and Ekins, 2020). These three compounds have therefore all made it to the clinic in various parts of the world for other applications (e.g. malaria, influenza, anti-protozoal etc.) and represent molecules that could be viable candidates for treating COVID-19. We now describe our efforts to repurpose these molecules for SARS-CoV-2.

**FIG 1.**
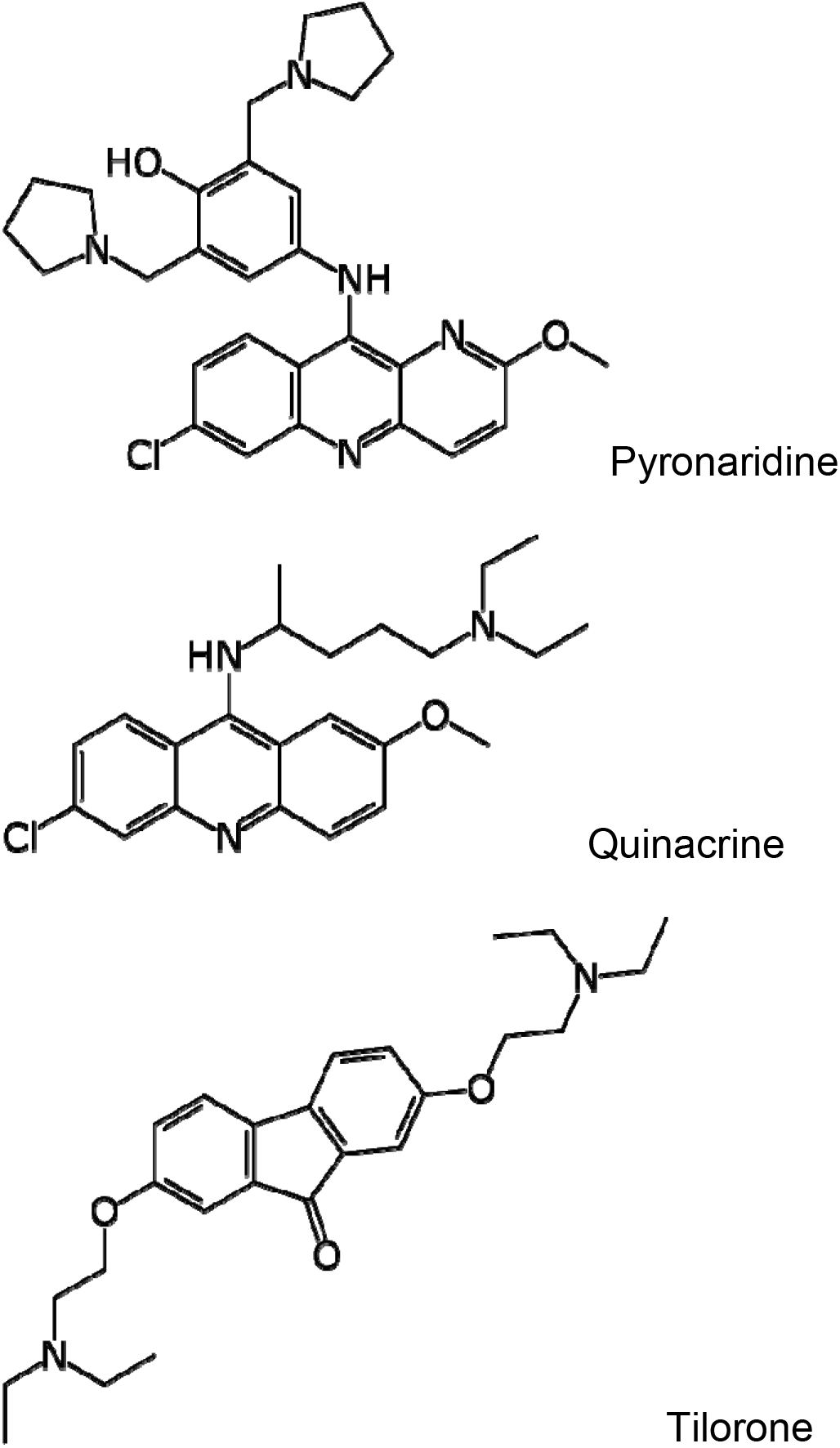
Structures of tilorone, quinacrine and tilorone.

## Materials and Methods

### Chemicals and reagents

Pyronaridine tetraphosphate [4-[(7-Chloro-2-methoxybenzo[b][1,5]naphthyridin-10-yl)amino]-2,6-bis(1-pyrrolidinylmethyl)phenol phosphate (1:4)] (Ekins et al., 2015a) was purchased from BOC Sciences (Shirley NY). Tilorone and quinacrine were purchased from Cayman Chemicals (Ann Arbor, Michigan). The purity of these compounds is greater than 95%.

#### Cell assays

##### Vero E6 cells

All drugs were tested with Vero E6 cells as described recently (Weston et al., 2020). Cells were plated in opaque 96-well plates 1 day prior to infection. Drug stocks were made in either dimethyl sulfoxide (DMSO), water, or methanol. Drugs were diluted from stock to 50⃼μM and an 8-point 1:2 dilution series made. Cells were pretreated with drug for 2⃼h at 37°C and 5% CO_2_ prior to infection at an MOI of 0.01 or 0.004. Vehicle controls were used on every plate, and all treatments were performed in triplicate for each screen. In addition to plates that were infected, parallel plates were left uninfected to monitor cytotoxicity of drug alone. Three independent screens with this setup were performed. Cells were incubated at 37°C and 5% CO_2_ for 3⃼days before CellTiter-Glo (CTG) assays were performed as per the manufacturer’s instructions (Promega). Luminescence was read using a Molecular Devices SpectraMax L plate reader.

##### A549-ACE2 cells

A549-ACE2 cells were plated in Corning black walled clear bottom 96 well plates 24 hours before infection for confluency. Drug stocks were diluted in DMSO for a 200X concentration in an 8-point 1:4 dilution series. Prepared 200X dilutions were then diluted to 2X concentration in infection media (Gibco DMEM supplemented with 5% HyClone FetalCloneII, 1% Gibco NEAA, 1% Gibco Pen-Strep). Growth media was removed and cells were pretreated with 2 X drug for 1 hour prior to infection at 37C and 5% CO2. Cells were either infected at a MOI of 0.02 with infectious clone SARS-CoV-2-nLuc (Hou et al., 2020) or mock infected with infection media to evaluate toxicity. 48 hours post infection wells were treated with Nano-Glo Luciferase assay activity to measure viral growth or CytoTox-Glo Cytotoxicity assay to evaluate toxicity of drug treatments, performed per manufacturer instructions (Promega). Nano-Glo assays were read using a Molecular Devices SpectraMax plate reader and CytoTox-Glo assays were read using a Promega GloMax plate reader. Vehicle treated wells on each plate were used to normalize replication and toxicity. Drug treatment was performed in technical duplicate and biological triplicate.

##### Vero 76 cells Reduction of virus-induced cytopathic effect (Primary CPE assay)

Confluent or near-confluent cell culture monolayers of Vero 76 cells are prepared in 96-well disposable microplates the day before testing. Cells are maintained in MEM supplemented with 5% FBS. For antiviral assays the same medium is used but with FBS reduced to 2% and supplemented with 50 μg/ml gentamicin. Compounds are dissolved in DMSO, saline or the diluent requested by the submitter. Less soluble compounds are vortexed, heated, and sonicated, and if they still do not go into solution are tested as colloidal suspensions. The test compound is prepared at four serial log_10_ concentrations, usually 0.1, 1.0, 10, and 100 μg/ml or μM (per sponsor preference). Lower concentrations are used when insufficient compound is supplied. Five microwells are used per dilution: three for infected cultures and two for uninfected toxicity cultures. Controls for the experiment consist of six microwells that are infected and not treated (virus controls) and six that are untreated and uninfected (cell controls) on every plate. A known active drug is tested in parallel as a positive control drug using the same method as is applied for test compounds. The positive control is tested with every test run.

Growth media is removed from the cells and the test compound is applied in 0.1 ml volume to wells at 2X concentration. Virus, normally at ~60 CCID_50_ (50% cell culture infectious dose) in 0.1 ml volume is added to the wells designated for virus infection. Medium devoid of virus is placed in toxicity control wells and cell control wells. Plates are incubated at 37 °C with 5% CO_2_ until marked CPE (>80% CPE for most virus strains) is observed in virus control wells. The plates are then stained with 0.011% neutral red for approximately two hours at 37°C in a 5% CO_2_ incubator. The neutral red medium is removed by complete aspiration, and the cells may be rinsed 1X with phosphate buffered solution (PBS) to remove residual dye. The PBS is completely removed, and the incorporated neutral red is eluted with 50% Sorensen’s citrate buffer/50% ethanol for at least 30 minutes. Neutral red dye penetrates living cells, thus, the more intense the red color, the larger the number of viable cells present in the wells. The dye content in each well is quantified using a spectrophotometer at 540 nm wavelength. The dye content in each set of wells is converted to a percentage of dye present in untreated control wells using a Microsoft Excel computer-based spreadsheet and normalized based on the virus control. The 50% effective (EC_50_, virus-inhibitory) concentrations and 50% cytotoxic (CC_50_, cell-inhibitory) concentrations are then calculated by regression analysis. The quotient of CC_50_ divided by EC_50_ gives the selectivity index (SI) value. Compounds showing SI values >10 are considered active.

##### Vero 76 cells Reduction of virus yield (Secondary VYR assay)

Active compounds are further tested in a confirmatory assay. This assay is set up like the methodology described above only eight half-log_10_ concentrations of inhibitor are tested for antiviral activity and cytotoxicity. After sufficient virus replication occurs (3 days for SARS-CoV-2), a sample of supernatant is taken from each infected well (three replicate wells are pooled) and tested immediately or held frozen at −80 °C for later virus titer determination. After maximum CPE is observed, the viable plates are stained with neutral red dye. The incorporated dye content is quantified as described above to generate the EC_50_ and CC_50_ values. The VYR test is a direct determination of how much the test compound inhibits virus replication. Virus yielded in the presence of test compound is titrated and compared to virus titers from the untreated virus controls. Samples were collected 3 days after infection. Titration of the viral samples (collected as described in the paragraph above) is performed by endpoint dilution (Reed and Muench, 1938). Serial 1/10 dilutions of virus are made and plated into 4 replicate wells containing fresh cell monolayers of Vero 76 cells. Plates are then incubated, and cells are scored for presence or absence of virus after distinct CPE is observed (3 days after infection), and the CCID_50_ calculated using the Reed-Muench method (Reed and Muench, 1938). The 90% (one log_10_) effective concentration (EC_90_) is calculated by regression analysis by plotting the log_10_ of the inhibitor concentration versus log_10_ of virus produced at each concentration. Dividing EC_90_ by the CC_50_ gives the SI value for this test.

##### Calu-3 cells

Calu-3 (ATCC, HTB-55) cells were pretreated with test compounds for 2 hours prior to continuous infection with SARS-CoV-2 (isolate USA WA1/2020) at a MOI=0.5. Forty-eight hours post-infection, cells were fixed, immunostained, and imaged by automated microscopy for infection (dsRNA+ cells/total cell number) and cell number. Sample well data was normalized to aggregated DMSO control wells and plotted versus drug concentration to determine the IC_50_ (infection: blue) and CC_50_ (toxicity: green).

##### Caco-2 cells

For the Caco-2 VYR assay, the methodology is identical to the Vero 76 cell assay other than the insufficient CPE is observed on Caco-2 cells to allow EC50 calculations. Supernatant from the Caco-2 cells are collected on day 3 post-infection and titrated on Vero 76 cells for virus titer as before.

##### Yield-reduction assays in monocytes, Calu-3 and Huh-7

Human hepatoma lineage (Huh-7), Lung epithelial cell line (Calu-3) or human primary monocytes from healthy donors (5 x10e5 cell/well in 24-multiwell plates) were infected at MOI of 0.1 for 1 h at 37 °C and treated with different concentrations of the compounds. Lysis of cell monolayer was performed 24 h (for monocytes) or 48 h (for Huh-7 and Calu-3 cells) post infection and culture supernatant was harvested 48 h post infection and virus was titrated by plaque-forming units (PFU) assays in Vero E6 cells. Alternatively, cell-associated viral genomic (ORF1b) and subgenomic (ORFE) RNA was quantified by real time RT-PCR (Wolfel et al., 2020). The standard curve method was employed for virus quantification. For reference to the cell amounts used, the housekeeping gene RNAse P was amplified (Sacramento et al., 2020).

##### VSV-pseudotype SARS-CoV-2 Neutralization Assays

Tilorone dihydrochloride, quinacrine hydrochloride, and pyronaridine tetraphosphate were tested for neutralization activity against the SARS-CoV-2 spike protein using a VSV-pseudotyped (rVSV-SARS-CoV-2 S) neutralization assay in Vero cells (IBT Bioscience (Rockville, MD)). The Luciferase-based microneutralization assay was conducted in Vero cells were seeded in black 96-well plates on Day1 at 6.00E+04 cells per well. Eight serial dilutions were prepared in triplicate and incubated for 1-hour with approximately 10,000 RLU of rVSV-SARS-CoV-2; virus only and cells only were added for controls and calculation. The TA/virus mixture was then added to the Vero cells and the plates were incubated for 24-hours at 37°C. Firefly Luciferase activity was detected using the Bright-Glo™ Assay System kit (Promega). Fifty percent inhibition concentration (IC_50_) was calculated using XLfit dose response model.

##### Murine Hepatitis Virus

Each compound was tested for antiviral activity against murine hepatitis virus (MHV), a group 2a betacoronavirus, in DBT cells. Each compound was tested against MHV using an 8-point dose response curve consisting of serial fourfold dilutions, starting at 10 μM. The same range of compound concentrations was also tested for cytotoxicity in uninfected cells.

##### HCoV 229E antiviral assay

HCoV 229E, (a gift from Ralph Baric, UNC, Chapel Hill) was propagated on Huh-7 cells and titers were determined by TCID_50_ assay on Huh-7 cells. Huh-7 cells were plated at a density of 25,000 cells per well in 96 well plates and incubated for 24 h at 37°C and 5% CO_2_. Cells were infected with HCoV 229E at a MOI of 0.1 in a volume of 50 ul MEM 1+1+1 (Modified Eagles Medium, 1% FBS, 1% antibiotics, 1% HEPES buffer) for 1 hour. Virus was removed, cells rinsed once with PBS and compounds were added at the indicated concentrations in a volume of 100 ul. Supernatants were harvested after 24 h, serially ten-fold diluted, and virus titer was determined by TCID_50_ assay on Huh-7 cells. CPE was monitored by visual inspection at 96h post infection. TCID_50_ titers were calculated using the Spearmann-Karber method (Kärber, 1931; Spearman, 1908).

##### Cytotoxicity

Cytotoxicity of compounds was assessed by MTT assay for quantification of cellular mitochondrial activity as an indirect measurement of cell viability. Briefly, freshly collected peripheral blood mononuclear cells (PBMCs) were plated in a 96 well plate at a concentration of 10⃼ cells/ well in RPMI medium for 2 h, to allow adhesion of monocytes. RPMI was then changed for complete medium supplemented with proper drug concentrations and controls for 24 h at 5% CO_2_ and 37°C. Vero CCL81 cells were cultivated at 5% CO_2_ and 37°C using Dulbecco’s Modified Eagle Medium supplemented with 10% heat-inactivated fetal bovine serum. For this experiment, Vero cells were seeded at a density of 10⃼ cells/ well in a 96 well plate prior incubation with a serial dilution of compounds of interest and controls for 72 h.

After drug treatment, cells were next incubated with 3-(4,5-Dimethylthiazol-2-yl)-2,5-Diphenyltetrazolium Bromide (Sigma-Aldrich M5655) for 4h followed by formazan crystal solubilization with isopropanol and absorbance readings at OD_570_ (Kumar et al., 2018). Cellular viability was expressed as a percentage relative to vehicle treated control. The CC_50_ was defined as the concentration that reduced the absorbance of treated cells to 50% when compared to non-treated controls. For Huh-7 cells, the cytotoxicity of extracts and pure compounds was determined according to the manufacturer's instructions using the CytoScan LDH cytotoxicity assay (G-Biosciences, St. Louis, MO). Briefly, 25,000 Huh-7 cells per well were added to 96 well plates and incubated for 24 h at 37°C and 5% CO_2_. Compounds were added with fresh media at the indicated concentrations to triplicate wells for 24h. Following the incubation, the plates were centrifuged at 250 x g for 5 min and 50 μl of supernatant from each well was transferred to a new plate. An equal volume of substrate mix was added to each well and the plates incubated at room temperature for 30 m. Then the stop solution was added, and the absorbance measured at 490 nm using a plate reader (Synergy HT, BioTek, Winooski, VT). Percent cytotoxicity was determined using the following formula: (Experimental-Spontaneous absorbance/Maximum-spontaneous absorbance) x 100.

##### Expression and purification of Spike RBD of SARS-CoV-2

A codon-optimized gene encoding for SARS-CoV-2 (331 to 528 amino acids, QIS60558.1) was expressed in Expi293 cells (Thermo Fisher Scientific) with human serum albumin secretion signal sequence and fusion tags (6xHistidine tag, Halo tag, and TwinStrep tag) as described before (Premkumar et al., 2020). S1 RBD was purified from the culture supernatant by nickel–nitrilotriacetic acid agarose (Qiagen), and purity was confirmed to by >95% as judged by coomassie stained SDS-PAGE. The purified RBD protein was buffer exchanged to 1x PBS prior to analysis by Microscale Thermophoresis.

##### Microscale Thermophoresis

Experiments were performed using a Monolith Pico (Nanotemper). Briefly, 10 μM protein was labelled using Monolith Protein Labeling Kit RED-NHS 2^nd^ Generation (Amine Reactive), with 3-fold excess NHS dye in PBS (pH 7.4). Free dye was removed according to manufacturer’s instruction, and protein was eluted in MST buffer (HEPES 10 mM pH 7.4, NaCl 150 mM), and centrifuged at 15 k rcf for 10 min. Binding affinity measurements were performed using 5 nM protein a serial dilution of compounds, starting at 250 μM. For each experimental compound, 16 independent stocks were made in DMSO using 2-fold serial dilution (10 mM initial concentration). 19.5 μL of Spike RBD (5 nM) of labeled protein in MST buffer containing 0.1% Triton X-100 and 1 mM BME was combined with 0.5 μL of the compound stock and then mixed thoroughly. This resulted in 2-fold serial dilution testing series with the highest and lowest concentration of 250 μM and 7.629 nM, respectively, with a consistent final DMSO concentration of 2.5%. Protein was incubated on ice in presence of compounds for one hour prior to transferring to standard Monolith NT.115 capillaries. Experiments were run at 20% excitation and high MST power at 23.0°C on a Monolith NT.115Pico (NanoTemper). Each experimental compound was run in triplicate.

The data were acquired with MO.Control 1.6.1 (NanoTemper Technologies). Recorded data were analyzed with MO.Affinity Analysis 2.3 (NanoTemper Technologies). The dissociation constant K_d_ quantifies the equilibrium of the reaction of the labelled molecule A (concentration c_A_) with its target T (concentration c_T_) to form the complex AT (concentration c_AT_): and is defined by the law of mass action as: 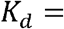 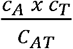, where all concentrations are “free” concentrations. During the titration experiments the concentration of the labelled molecule A is kept constant and the concentration of added target T is increased. These concentrations are known and can be used to calculate the dissociation constant. The free concentration of the labelled molecule A is the added concentration minus the concentration of formed complex AT. The K_d_ is calculated as 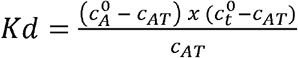. The fraction of bound molecules x can be derived from F_norm_, where F_norm_(A) is the normalized fluorescence of only unbound labelled molecules A and F_norm_(AT) is the normalized fluorescence of complexes AT of labeled as shown by the equation: 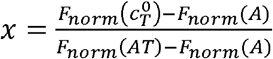. The MST traces that showed aggregation or outliers were removed from the datasets prior to Kd determination.

## Results

### Cell assays

SARS-CoV-2 susceptibility to tilorone, quinacrine and pyronaridine was determined in two lineages of Vero Cells for initial screening. For Vero E6 (Fig. S1) and 76 cell lines (Table 1), tilorone emerged as a potential hit, because of 7.5-fold margin between cytotoxicity and potency, as judged by the selectivity index (SI) (Table 1). Moving forward to determine if these compounds were endowed with anti-SARS-CoV-2 activity in cells relevant for COVID-19 pathophysiology, antiviral activity was assayed in intestinal, respiratory, hepatic and immune cells. Testing in a VYR assay on Caco-2 cells demonstrated activity for all three molecules (Table 2) with quinacrine EC_90_ 10.54 μM (CC_50_ 229.15 μM), tilorone EC_90_ 28.96 μM (CC_50_ 111.49) and pyronaridine EC_90_ 5.49 μM (CC_50_ 51.65 μM).

**Table 1.**
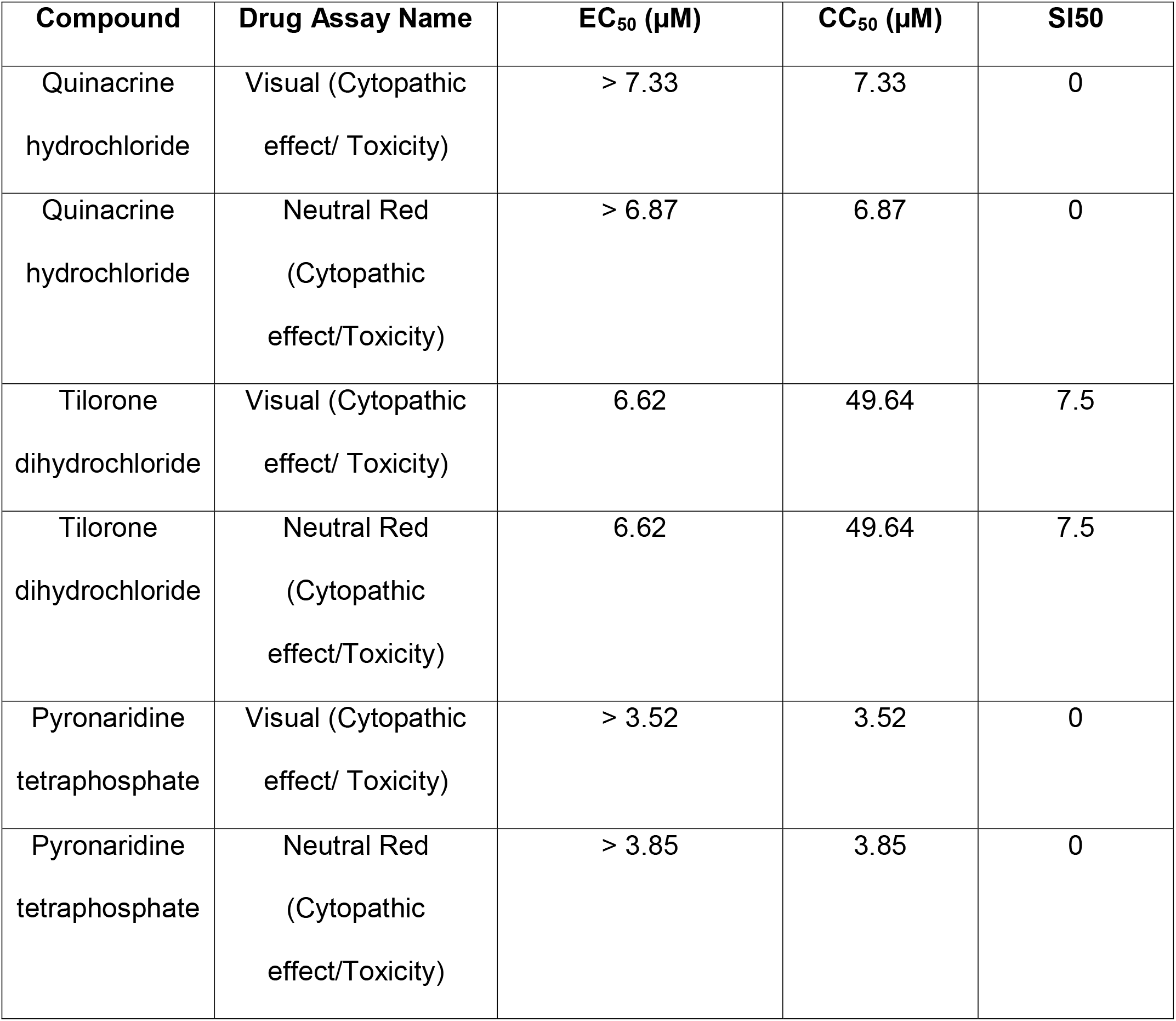
EC_50_ and CC_50_ values for quinacrine, pyronaridine and tilorone against SARS-CoV-2 (strain USA_WA1/2020) in Vero 76 cells. Drug concentration range: 0.1-100 μg/mL.

| **Compound** | **Drug Assay Name** | **EC<sub>50</sub> (µM)** | **CC<sub>50</sub> (µM)** | **SI<sub>50</sub>** |
| --- | --- | --- | --- | --- |
| Quinacrine hydrochloride | Visual (Cytopathic effect/ Toxicity) | > 7.33 | 7.33 | 0 |
| Quinacrine hydrochloride | Neutral Red (Cytopathic effect/Toxicity) | > 6.87 | 6.87 | 0 |
| Tilorone dihydrochloride | Visual (Cytopathic effect/ Toxicity) | 6.62 | 49.64 | 7.5 |
| Tilorone dihydrochloride | Neutral Red (Cytopathic effect/Toxicity) | 6.62 | 49.64 | 7.5 |
| Pyronaridine tetraphosphate | Visual (Cytopathic effect/ Toxicity) | > 3.52 | 3.52 | 0 |
| Pyronaridine tetraphosphate | Neutral Red (Cytopathic effect/Toxicity) | > 3.85 | 3.85 | 0 |

**Table 2.**
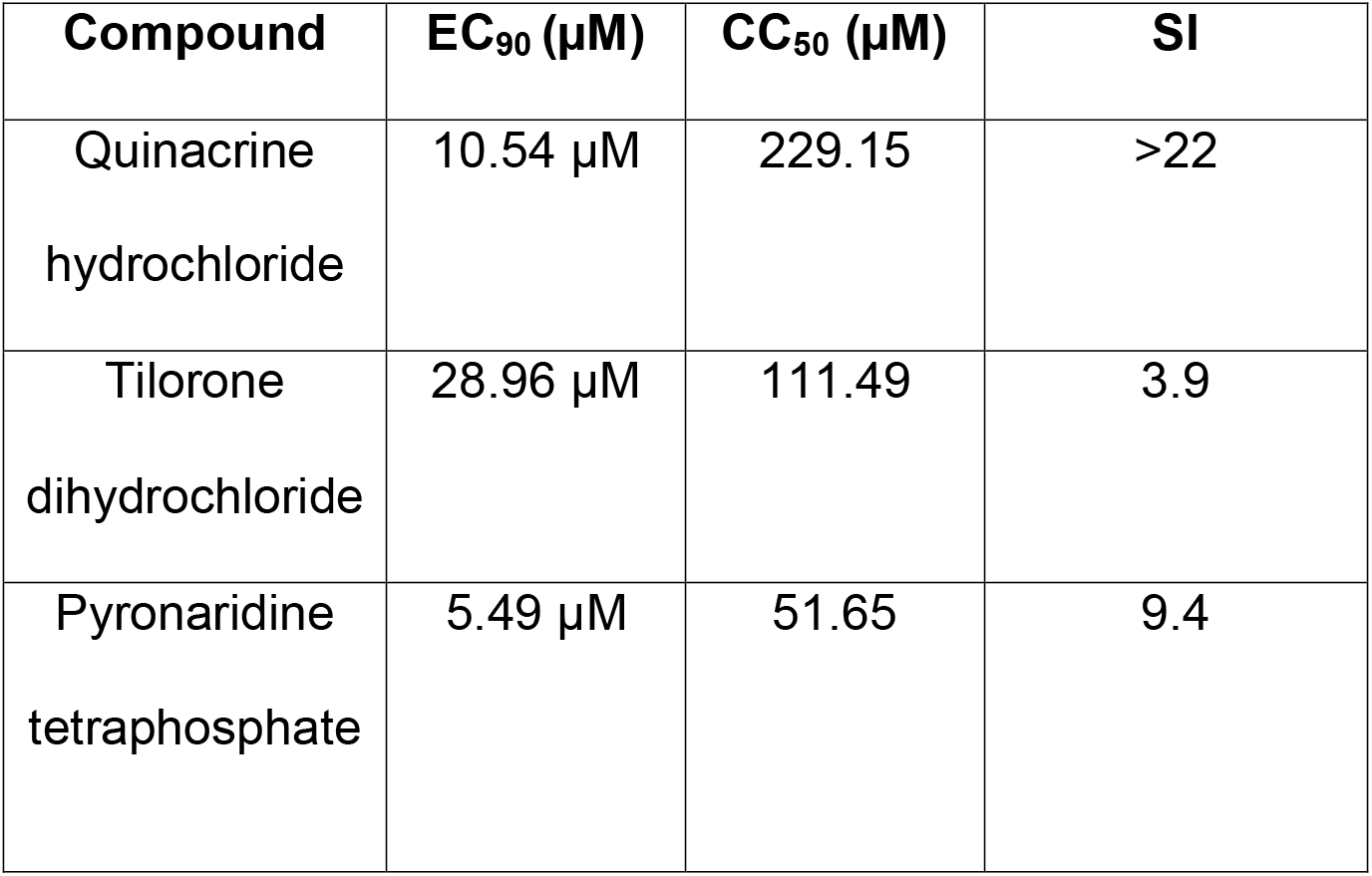
EC_90_ and CC_50_ values for Quinacrine and Tilorone against SARS-CoV-2 (strain USA_WA1/2020) in Caco-2 cells. Drug concentration range: 0.032-100 μg/mL. No CPE observed in this assay. Only VYR data was reported.

| **Compound** | **EC<sub>90</sub> (µM)** | **CC<sub>50</sub> (µM)** | **SI** |
| --- | --- | --- | --- |
| Quinacrine<br>hydrochloride | 10.54 µM | 229.15 | >22 |
| Tilorone<br>dihydrochloride | 28.96 µM | 111.49 | 3.9 |
| Pyronaridine<br>tetraphosphate | 5.49 µM | 51.65 | 9.4 |

By means of measuring the double-stranded virus RNA as a proxy of virus replication, tilorone showed activity in Calu-3 cells with an EC_50_ of 10.77 μM, and SI=2 (Fig. S2). For comparison, remdesivir showed an EC_50_ of 0.016 μM and an SI>622 (Fig. S2). Moreover, when supernatant from SARS-CoV-2-infected Calu-3 cells treated with tilorone was collected and assay for its ability to perform another round of infection in highly permissive VeroE6 cells, 80 ± 10 % inhibition of virus production was quantified at 1 μM (Fig. S3). At this same concentration, remdesivir inhibited 99 ± 1 % virus production (Fig. S3).

Tilorone was also tested in another laboratory (Dr. Thiago Moreno, Fiocruz, Brazil) at MOI of 0.1 for 1 h at 37 °C and at different concentrations. Lysis of cell monolayer was performed 48 h post infection and virus was titrated by plaque-forming units (PFU) assays and reported as % inhibition. In this assay, tilorone had an IC_50_ ~ 9 μM and remdesivir showed almost 100 % inhibition PFU/mL even at the lowest concentration tested 0.6 μM (Fig. S4, reported as % inhibition (C) and PFU/mL (D)).

SARS-CoV-2 can infect Human hepatoma lineage (Huh-7), however, the viral titers produced in this cell are lower than in Vero and Calu-3 (Sacramento et al., 2020). Whereas African green monkey kidney cell (Vero E6), human hepatoma (HuH-7) and airway epithelial cells (Calu-3)(Yoshikawa et al., 2009) produced infectious SARS-CoV-2 titers and quantifiable RNA levels, A549 pneumocytes and induced pluripotent human neural stem cells (NSC) displayed limited ability to generate virus progeny, as measured by plaque forming units (PFU) of virus bellow the limit of detection. In Human hepatoma lineage (Huh-7), these compounds did not show antiviral activity up to 1 μM (Fig. S3 A).

Tilorone was found to impair SARS-CoV-2 replication in human primary monocytes at 10 μM. Human primary monocytes were infected at the indicated MOI of 0.1 and treated with 10 μM of tilorone or remdesivir for 24 h. Tilorone showed similar inhibition of SARS-CoV-2 measured by the cell-associated viral RNA levels quantified by real time RT-PCR, compared to remdesivir when cells were treated with 10 μM of compounds (Fig. S3B). MTT Cytotoxicity assays in human monocytes and Vero CCL81 cells demonstrated similar CC_50_ for quinacrine and pyronaridine (Fig. S4).

The most promising results by far were achieved in A549-ACE2 cells which support SARS-CoV2 growth to about 10^7^ pfu/ml and in which quinacrine, tilorone and pyronaridine all showed SARS-CoV-2 inhibition demonstrating IC_50_ values < 200 nM (Fig. 2) and good selectivity indices. This inhibition compares well with remdesivir under the same conditions.

**FIG 2.**
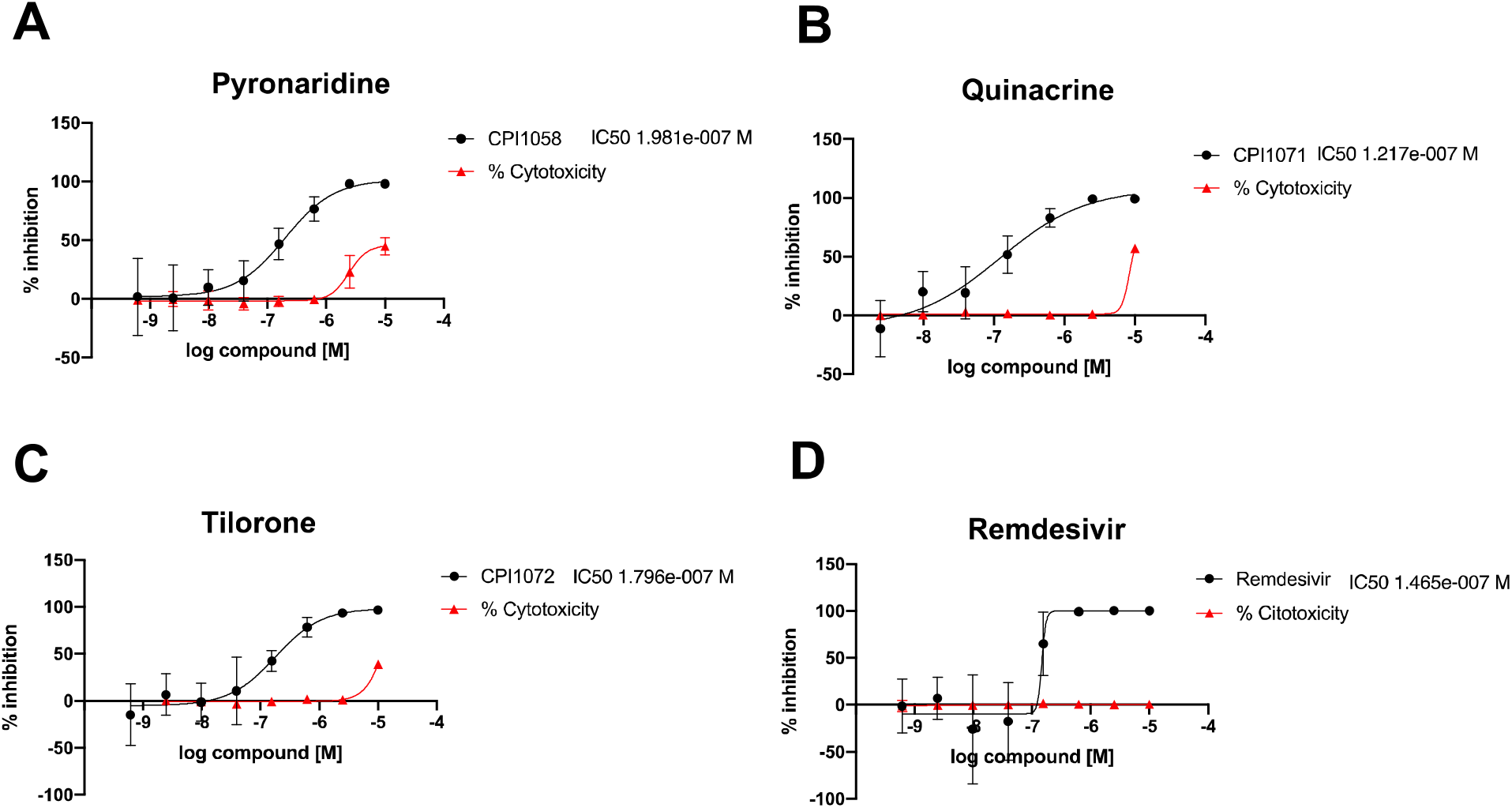
SARS-CoV-2 inhibition in A549-ACE2 cell lines. A) Pyronaridine IC_50_ = 198 nM, B) Quinacrine IC_50_ = 122 nM, C) Tilorone IC_50_ = 180 nM and D) Remdesivir IC_50_ = 147 nM.

All three compounds were also tested against a group 2a murine hepatitis virus (MHV), in DBT cells, a model of betacoronavirus genetics and replication (Yount et al., 2002). Quinacrine showed an IC_50_ 2.3 μM, pyronaridine showed an IC_50_ 2.75 μM while for tilorone the dose response curve did not reach the plateau and the IC_50_ was estimated to be 20 μM (Fig. 3).

**FIG 3.**
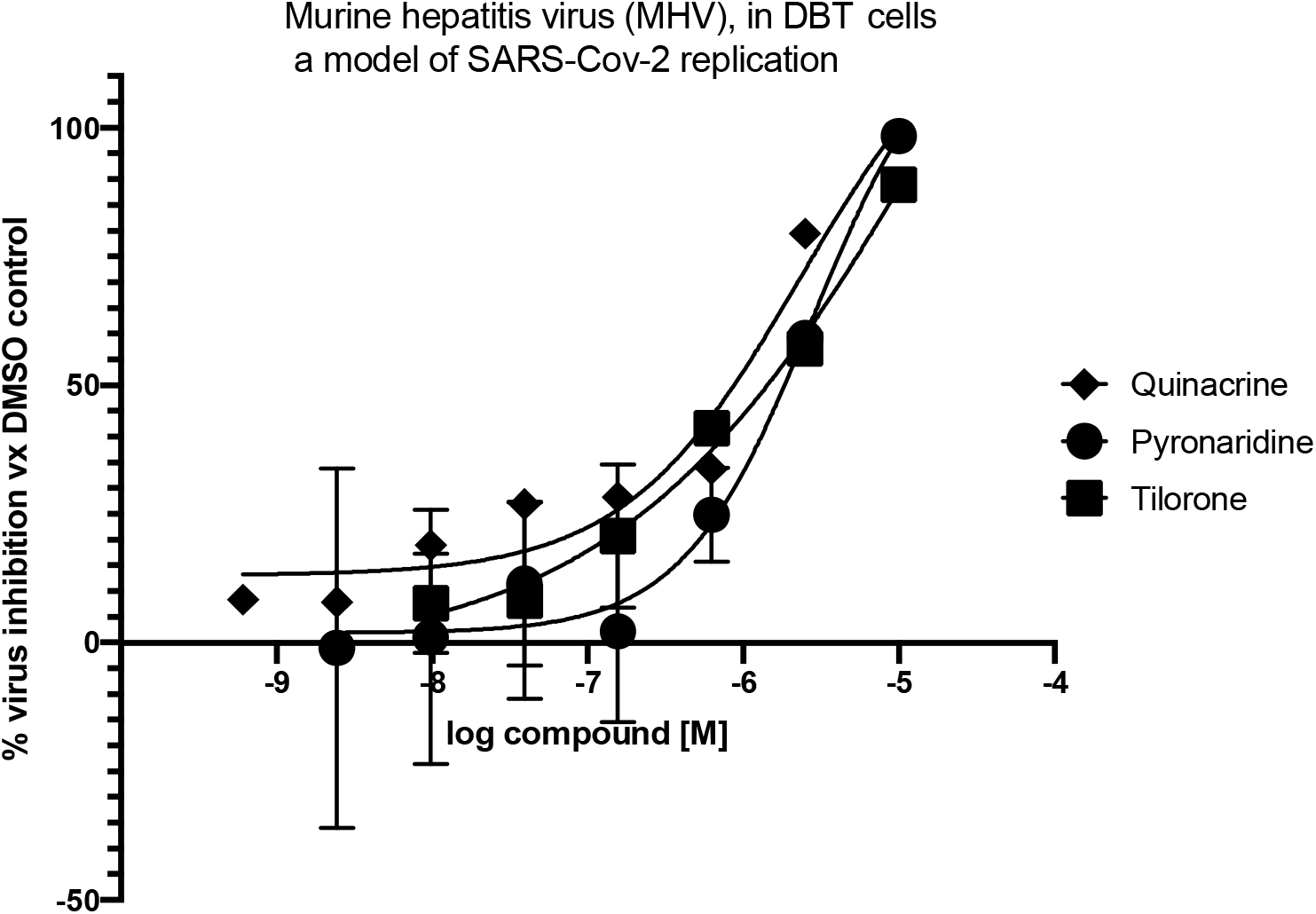
Murine hepatitis virus (MHV), in DBT cells, a model of SARS-Cov-2 replication. Quinacrine IC_50_ = 2.3 μM, Pyronaridine IC_50_ = 2.75 μM. Tilorone dose response curve did not reach the plateau and the IC_50_ was estimated to be 20 μM.

These compounds were also tested in Huh-7 cells infected by the human coronavirus 229E (HCov-229E), a group A alphacoronavirus which infects humans and bats (Corman et al., 2016; Lim et al., 2016). It is an enveloped, positive-sense, single-stranded RNA virus which enters its host cell by binding to the aminopeptidase N (AP-N) receptor (Fehr and Perlman, 2015). Quinacrine showed a decrease of 3.9 logTCID50/ml when tested at 10 μM, pyronaridine showed a decrease of 2.83 logTCID50/ml when tested at 20 μM and tilorone did not show significant inhibition. The CC_50_ was > 15 μM for quinacrine and CC_50_ was > 20 μM for pyronaridine (Fig. S5).

### Microscale Thermophoresis

Based on our previous work showing that pyronaridine, tilorone and quinacrine bind to the EBOV glycoprotein (Lane and Ekins, 2020), this provided impetus to test them against the SARS-CoV-2 Spike RBD. The K_d_ values for tilorone and pyronaridine were of 339 nM and 647 nM, respectively at pH 7.4 (Fig. 4A) and K_d_ 631nM and 618nM at pH5.2, respectively (Fig. 4B). Quinacrine did not demonstrate reproducible binding to this protein.

**FIG 4.**
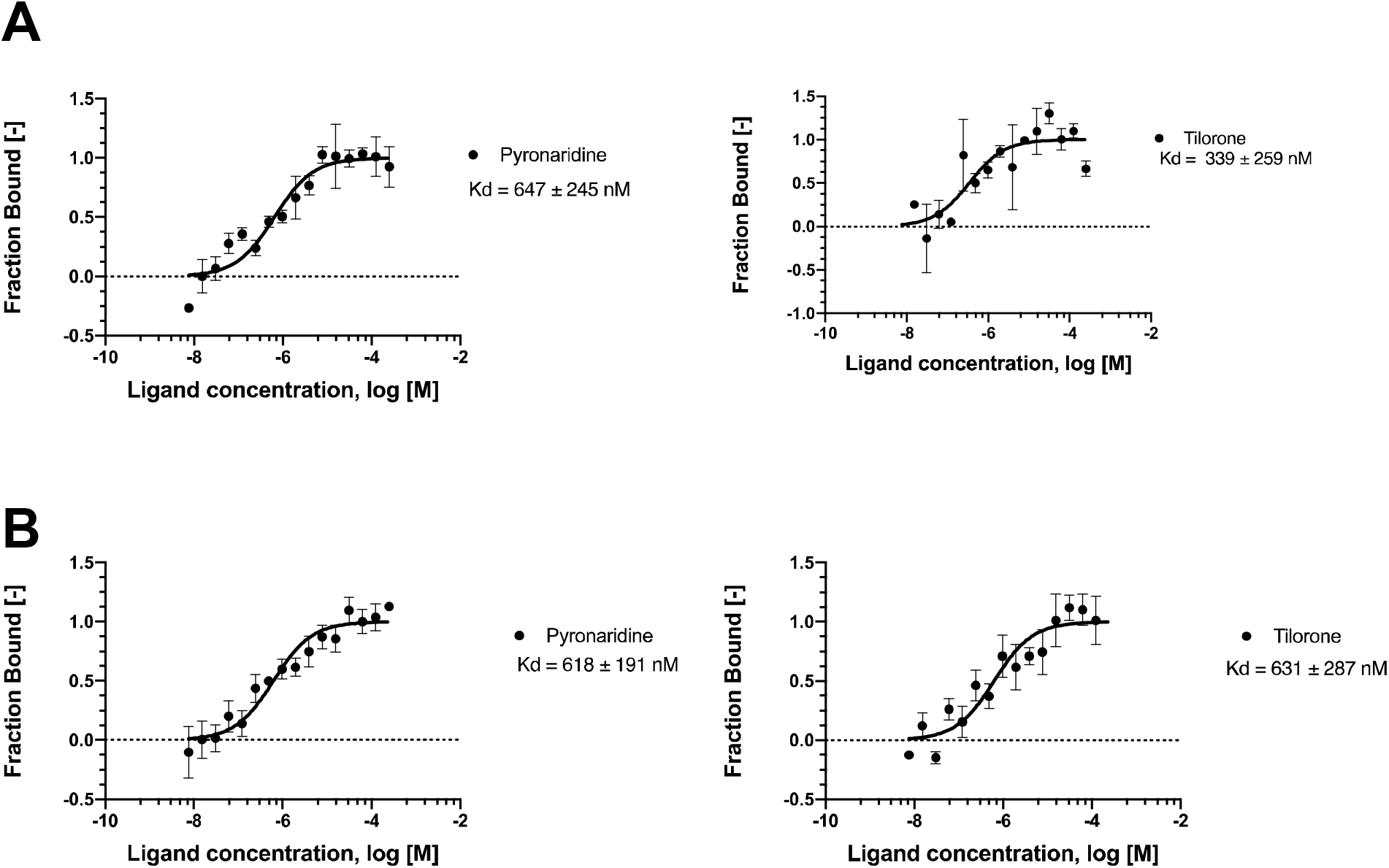
MicroScale Thermophoresis binding analysis for the interaction between Spike RBD and compounds. The concentration of labeled Spike RBD is kept constant at 5 nM, while the ligand concentration varies from 250 μM and 7.629 nM. The serial titrations result in measurable changes in the fluorescence signal within a temperature gradient that can be used to calculate the dissociation constant. The curve is shown as Fraction Bound [-] against compound concentration on a log scale. Binding affinity was measured at pH 7.4. **(A)** a pH 5.2 **(B)**.

### VSV-pseudotype SARS-CoV-2 Neutralization Assays

To directly measure the impact of these drugs on SARS-CoV2 S glycoprotein mediated entry, we employed VSV-pseudotype SARS-CoV2 assays. The neutralization activity for tilorone and pyronaridine did not achieve 50% neutralization as the dose response curves did not reach the plateau, suggesting little if any activity against SARS-CoV-2 S glycoprotein mediated docking and entry (Fig. S6 and Table S2).

## Discussion

Identifying drugs for any new virus in real time is extremely challenging due to the pressure to identify a treatment while large numbers of patients are suffering or dying, with only palliative care available. The SARS-CoV-2 outbreak is only the most recent such example. In humans, SARS-CoV-2 is currently thought to cause a biphasic disease characterized by early high titer virus replication in airway epithelial cells and type II pneumocytes, followed by virus clearance and immune mediated pathology (Gan et al., 2020). Consequently, drug development against SARS-CoV-2 is complicated by the diverse disease mediating mechanisms associated with early direct virus cell killing and late immune mediated pathology (Gan et al., 2020). Studies in animals and in humans demonstrate that early administration of direct acting antivirals is essential for efficacy, however, later in infection combination therapies including direct acting antivirals with anti-inflammatory drugs will likely be required. Most of the research emphasis to date has been on the development of vaccines or biologics and only a relatively few small molecules have made it to the clinic such as remdesivir (Beigel et al., 2020a, b). Remdesivir was originally developed for HCV then repurposed for EBOV, therefore we hypothesized that other drugs used for treating this latter virus should also be evaluated. Our focus on repurposing drugs for EBOV (Ekins et al., 2015a) previously has led us to apply our approaches to SARS-CoV-2 (Gawriljuk et al., 2020). We reasoned that the three molecules (pyronaridine, tilorone and quinacrine) for which we already had some antiviral knowledge of may be a useful starting point to explore for repurposing for SARS-CoV-2.

Several groups have now published on these three drugs and in part due to these efforts to raise visibility, pyronaridine (Anon, 2020a) and tilorone (Anon, 2020b) are in clinical trials in different countries. Tilorone was previously identified *in vitro* in Vero cells as a hit against SARS-CoV-2 (IC_50_ 4 μM) (Jeon et al., 2020b) and has also been shown to have similar activity against MERS (Ekins and Madrid, 2020a) and Ebola (Ekins et al., 2015b) (as has remdesivir (de Wit et al., 2020)). Tilorone was recently found to block the endocytosis of preformed α-Syn fibrils and was an inhibitor of HSPG-dependent endocytosis (Zhang et al., 2020). Combined with our earlier findings of tilorone binding to EBOV glycoprotein (Lane and Ekins, 2020), this would suggest a direct antiviral effect rather than an effect on the innate immune system, (as long assumed since the 1970s)(Ekins et al., 2020), is responsible for the efficacy observed in this case. In our hands tilorone showed antiviral activity against SARS-CoV-2 in A549-ACE2 (IC_50_ 180 nM), Vero 76 cell lines (IC_50_ of 6.62 μM) but not Vero E6, and to a lesser extent in Caco-2, Calu-3, and monocytes as well.

Pyronaridine is used in combination with artesunate to treat malaria (Croft et al., 2012). We have previously demonstrated that both molecules show additivity against EBOV (Lane et al., 2020a). Pyronaridine was therefore of interest to us as a potential antiviral against SARS-CoV-2. A preprint of the Jeon et al., paper (Jeon et al., 2020b) included pyronaridine (IC_50_ 31 μM) but this was removed before publication. More recently pyronaridine was tested again in Vero cells by others (IC_50_ 1.08 μM, CC_50_ 37.09 μM) at 24 hpi and in Calu-3 cells (IC_50_ 6.4 μM CC_50_ 43.08 μM) at 24 hpi (Bae et al., 2020). Our results do not match any of this earlier data as we showed no activity in Vero 76, Vero E6 cells, Calu-3 cells, while we did see activity in Caco-2 (EC_90_ 5.49 μM), A549-ACE2 cells (IC_50_= 198 nM) and MHV (IC_50_ 2.75 μM).

A recent screen of the Prestwick library in hPSC lung organoids identified the antiprotozoal quinacrine (EC_50_ 2.83 μM) (Han et al., 2020) which was followed up in mice infected with SARS-CoV-2 pseudovirus and showed a significant decrease in infected cells (Han et al., 2020). Others have not observed significant *in vitro* SARS-CoV-2 activity for quinacrine in Vero cells (Jeon et al., 2020b) while it has previously been demonstrated to possess activity against Ebola infected HeLa cells (Ekins et al., 2015b) but not Vero cells (Lane et al., 2019a). In this study we confirm no *in vitro* activity in Vero E6, Vero76, Calu-3 and demonstrate activity in Caco-2 (EC_90_ 10.54 μM), A549-ACE2 (IC_50_ 122 nM) and MHV (IC_50_ 2.3 μM) for quinacrine.

These three drugs have therefore shown considerable variability in testing in different cell types infected with SARS-CoV-2 (Table S1) compared to recent literature. The differences between these and previous data published in Vero cells could be due to a number of factors including differences in: assays, MOI, time of addition of the drug and expression levels of ACE2. It should also be clear that we see differences in the data reported for these compounds as well. These three drugs have all demonstrated low μM activity against SARS-CoV-2 in the best cases, while A549-ACE2 seems especially sensitive to these drugs and the remdesivir data is in line with published data in different cell lines (Pruijssers et al., 2020). Our observations of no inhibition in Vero E6 cells for tilorone, quinacrine and pyronaridine is exactly as we had observed previously for EBOV (Lane et al., 2019b). The gold standard currently are primary human airway epithelial cells or a primary type II ATII cell as are targeted by the virus *in vivo*.

We characterized binding of pyronaridine and tilorone to the Spike RBD using MST, with K_d_ values for tilorone and pyronaridine of 339 nM and 647 nM, respectively (Fig. 4). Pyronaridine and tilorone bind to the Spike RBD with 20-40 times weaker affinity when compared to ACE2, which has been reported to be ~ 15 nM by different techniques (Chan et al., 2020; Wrapp et al., 2020). In the VSV-pseudotype SARS-CoV-2 neutralization assay we saw no measurable activity for tilorone and pyronaridine (Fig. S6 and Table S2) suggesting that it may be necessary to modify these compounds to increase affinity for the spike protein to enhance activity. The binding affinity experiments using MST were performed with the RBD, which is the receptor binding domain that binds to ACE2. Despite these compounds binding to the Spike RBD, their affinity is clearly not high enough to compete with binding to ACE2 in the pseudovirus assay, or the compounds may bind to the RBD in a place that does not affect binding to the ACE2 receptor. We have previously characterized that these compounds are lysosomotropic (Lane et al., 2020a) so this may also be their mechanism of action against SARS-CoV-2.

The effect of lysosomotropic compounds on cells is multifaceted as evidenced by the prototypical lysosomotropic compound chloroquine. A well-known antiviral effect of chloroquine is against EBOV and has been associated with the pH increase within lysosomes, which reduces the efficiency of acid hydrolases (Cathepsins) required for viral glycoprotein priming. As there is some evidence that cathepsin is also involved in one entry mechanism of SARS-CoV-2, this function may also be involved in the partial inhibition of SARS-CoV-2 by compounds of this class although it is unlikely the only mechanism of inhibition by lysosomotropic compounds. Interestingly, SARS-CoV-2 has recently been shown to also increase the pH of lysosomes, possibly through the open reading frame protein 3A (ORF3a) (Ghosh et al., 2020). As the Sarbecovirus E protein also functions as a viroporin, these overlapping activities are likely requisites for efficient SARS-CoV-2 infection (Castano-Rodriguez et al., 2018; Lu et al., 2006; Yount et al., 2005; Yue et al., 2018). ORF3a has been shown to traffic to lysosomes and disrupt their acidification and ultimately viral egress (Ghosh et al., 2020). ORF3a is a viroporin, which modifies several cellular functions, including membrane permeability, membrane remodeling and glycoprotein trafficking (Nieva et al., 2012). While lysosomotropic compounds have been shown to increase lysosomal pH this effect has been demonstrated to be transient with multiple lysosomotropic compounds including chloroquine, fluoxetine, tamoxifen and chloropromazine. This was illustrated by multiple types of measurements, including a lack of a decrease in cathepsin activity over time (Lu et al., 2017). It is possible that since cells are able to counter the pH change caused by the lysosomotropic compounds, this may translate into this same phenomenon with SARS-CoV-2 infected cells. If an increased pH of the lysosome is indeed required for the efficient egress of the virus, compounds that counteract this would potentially act as an antiviral. The “redundant” viroporin activities of E protein and ORF3a may also complicate the specificity and potency of lysosomotropic compounds if they target one, but not both proteins. This is consistent with the ability to delete either one separately and recover viable viruses, yet if both are deleted, then the virus is inhibited. Consequently, lysosomotropic inhibitors must target both activities.

While a pH change in acidic organelles is the most well-known effect of lysosomotropic compounds, they are also known to elicit other cellular effects such as inducing vacuole formation (Marceau et al., 2012) (Mauthe et al., 2018) by the fusion of lysosomes and late endosomes (Kaufmann and Krise, 2008). This is an important distinction because a precursor to this is a general disorganization of the Golgi complexes with an increased number of vesicles found proximal to the Golgi (Mauthe et al., 2018). Therefore, lysosomotropic compounds not only create vacuoles, but also cause the disruption of vesicle translation from the Golgi to distal subcellular locations (Chen et al., 2011; Kellokumpu et al., 2002; Rivinoja et al., 2009). Many lysosomotropic compounds have shown a similar disruption in vesicle trafficking, including transport to the apical plasma membrane (Ellis and Weisz, 2006; Matlin, 1986). Disruption of this translocation of vesicles from the TGN would likely inhibit the common biosynthetic secretory pathway used for egress of multiple viruses such as hepatitis C virus, dengue virus, and West Nile virus (Ravindran et al., 2016; Robinson et al., 2018). Additionally, many drugs that are known to be lysosomotropic also induce phospholipidosis in cells (Orogo et al., 2012) which is the reduced degradation of phospholipids, causing an excess accumulation in cells. The mechanism of drug-induced phospholipidosis is unclear but could be due to the cationic drug binding to bis(monoacylglycero)phosphate (BMP) in the phospholipid bilayer, which is heavily enriched within the lysosome membrane (Shayman and Abe, 2013). This affects the efficiency of acid hydrolases as well as disrupts interactions between membrane-bound proteins and molecules within the lysosomal lumen. Drug-induced phospholipidosis is also suggested to alter the lysosomal membrane curvature (Baciu et al., 2006) similar to pH (Lahdesmaki et al., 2010). A change in membrane curvature is a phenomenon suggested to be important for the membrane fusion of other viruses (Stiasny and Heinz, 2004), which points to a similar effect on the membrane fusion of SARS-CoV-2. Multiple cellular alterations by lysosomotropic compounds may be involved in the inhibition of SARS-CoV-2 egress.

The relevance of pyronaridine, tilorone and quinacrine as antivirals can be further assessed using their concentrations attained *in vivo*. The C_max_ data in our previous mice pharmacokinetics studies (i.p. dosing) suggests the concentration is above the average IC_50_ observed for EBOV inhibition *in vitro* (~1 μM) (Table S2). For quinacrine and pyronaridine we have also identified published human pharmacokinetics data and these suggest that IC_50_ values up to 1 μM would be below the C_max_ achieved for quinacrine and pyronaridine (Table S3). Therefore, as we have demonstrated IC_50_ values in some cell lines infected with SARS-CoV-2 around or below 1 μM which may enable them to achieve efficacious concentrations. While we could not find human pharmacokinetics data for quinacrine it can still be considered as it was safely used during WWII in millions of soldiers as an antimalarial (Lane et al., 2019a). These three molecules generally have excellent *in vitro* ADME properties and there is a considerable body of data we have built up on them (Table S4) including maximum tolerated dose in mice. This would certainly be useful for designing efficacy assessment studies in SARS-CoV-2 infected mouse models in future.

It is important to study virus infection in different cell lines and understand the subtle differences among them when treated with drugs. SARS-CoV-2 infection experiments using primary human airway epithelial cells have been found to have cytopathic effects 96 h after the infection (Zhu et al., 2020). However, primary human airway epithelial cells are expensive and do not proliferate indefinitely (Takayama, 2020). Several infinitely proliferating cell lines, such as Caco-2 (Kim et al., 2020), Calu-3 (Ou et al., 2020), HEK293T (Harcourt et al., 2020), and Huh7 (Ou et al., 2020) have been utilized in SARS-CoV-2 infection experiments. These cell lines do not accurately mimic human physiological conditions and generate low titers of infectious SARS-CoV-2 (Harcourt et al., 2020; Kim et al., 2020; Ou et al., 2020). Despite this limitation, valuable information about the virus infection and replication can be learned from studies using these cell lines. A previous study (Chu et al., 2020) assessed 25 cell lines derived from different tissues or organs and host species and reported that cytopathic effects were only seen in VeroE6 and FRhK4 cells after SARS-CoV-2 inoculation for up to 120 hpi. These findings are important for optimization of antiviral assays based on cell protection assessment, because cell lines without obvious cytopathic effects might lead to overestimation of cell viability and drug efficacy (Chu et al., 2020). For efficient SARS-CoV-2 research, a cell line, such as Vero cells, that can easily replicate and isolate the virus is essential, but they have been shown not to produce interferon (IFN) when infected with Newcastle disease virus, rubella virus, and other viruses (Desmyter et al., 1968). Under previously described experimental conditions, productive SARS-CoV-2 replication in A549 cells was erratic (Sacramento et al., 2020) which can be overcome by preparing A549 cells overexpressing ACE2 (as used in the current study).

The differences in responses in different cell lines could be accounted for by the basic biochemistry, for example hepatic cells, like Huh-7, are equipped with enzymes to synthesize nucleotides, carbohydrates and lipids (Nwosu et al., 2018). Hence it is not surprising that the highest potency of remdesivir against SARS-CoV-2 is found in Huh-7 cells, followed by Calu-3 and Vero, meaning that Huh-7, and subsequently, Calu-3 are active to convert it to its triphosphate (Dittmar et al., 2020; Sacramento et al., 2020).

While nasal airway epithelium replicates virus best early (Hou et al., 2020) Type II pneumocytes are the most affected cell in the lung of patients that died from COVID-19 (Carsana et al., 2020). Both A549 and Calu-3 cells are lung epithelial cells (based on ATCC information). Under continuous submersible culture A549 cells decrease the expression of type C surfactant and enhances the expression aquaporin V, a phenotype of type I pneumocytes (Wu et al., 2017). Calu-3 is an airway epithelial cell that can be induced to differentiate into a ciliated airway epithelial cell (Yoshikawa et al., 2009). For entry inhibitors, Calu-3 is a better model than Vero and A549 cells, because Calu-3 expresses TMPRSS2. The lack of antiviral activity of chloroquine in Calu-3 cells would likely have anticipated its failure in clinical trials (Hoffmann et al., 2020b). Under regular cell culture Calu-3 and Caco-2 better support virus entry than A549 (Chu et al., 2020; Hoffmann et al., 2020a) however the latter are much easier to grow, reinforcing the interest in generating A549-ACE2 cells which replicate SARS-CoV2 to titers of ~10^7^ PFU/ml.

In conclusion, this study shows the importance of an exhaustive comparison of different cell lines when testing small molecules as inhibitors of SARS-CoV-2 and clearly indicates how subtle differences in experimental approaches with the same cell lines can have dramatic effects on whether a drug is identified as an inhibitor or not. We illustrate this now for Vero cells which when infected with Ebola or SARS-CoV-2 are insensitive to quinacrine, tilorone and pyronaridine (Lane et al., 2019b). This may be related to the mechanism of entry in these cells, the lack of IFN in Vero cells or other factors that limits activity for these lysosomotropic compounds (Lane et al., 2020a). While they are not as potent inhibitors against SARS-CoV-2 as remdesivir in most cell types, they are comparable in the A549-ACE2 cell line. Future work will focus on testing the efficacy of these drugs against SARS-CoV-2 in animal models and further interrogation of mechanism.

## Acknowledgments

Victor O. Gawriljuk is kindly acknowledged for helping collate literature data on SARS-CoV-2. We graciously thank Dr. Sara Cherry and Dr. David Schultz for the Calu-3 high-content SARS-CoV-2 studies performed by the University of Pennsylvania High-throughput Screening Core and the Cherry laboratory.

Dr. Mindy Davis and colleagues are gratefully acknowledged for assistance with the NIAID virus screening capabilities.

## Funding

We kindly acknowledge NIH funding: R44GM122196-02A1 from NIGMS (PI – Sean Ekins), 1R43AT010585-01 from NIH/NCCAM, AI142759 and AI108197 to RSB, and support from DARPA (HR0011-19-C-0108; PI: P. Madrid) is gratefully acknowledged. FTMC and TAT are supported by FAPESP (grant 2017/18611-7 and grant 2020/05369-6 for FTMC, and grant 2019/27626-3 for TAT). Distribution Statement “A” (Approved for Public Release, Distribution Unlimited). The views, opinions, and/or findings expressed are those of the author and should not be interpreted as representing the official views or policies of the Department of Defense or the U.S. Government. This project was also supported by the North Carolina Policy Collaboratory at the University of North Carolina at Chapel Hill with funding from the North Carolina Coronavirus Relief Fund established and appropriated by the North Carolina General Assembly.

Collaborations Pharmaceuticals, Inc. has utilized the non-clinical and pre-clinical services program offered by the National Institute of Allergy and Infectious Diseases.

