## Supplemental data files for "Repurposing the Ebola and Marburg Virus Inhibitors Tilorone, Quinacrine and Pyronaridine: In vitro Activity Against SARS-CoV-2 and Potential Mechanisms"

**Short running title:** Ebola SARS-CoV-2 inhibitors

**Authors:** Ana C. Puhl^a#^, Ethan James Fritch^b^, Thomas R. Lane^a^, Longping V. Tse^c^, Boyd L. Yount^c^, Carol Queiroz Sacramento^d,e^, Tatyana Almeida Tavella^f^, Fabio Trindade Maranhão Costa^f^, Stuart Weston^g^, James Logue^g^, Matthew Frieman^g^, Lakshmanane Premkumar^b^, Kenneth H. Pearce^h,i^, Brett L. Hurst^j,k^, Carolina Horta Andrade^f,l^, James A. Levi^m^, Nicole J. Johnson^m^, Samantha C. Kisthardt^m^, Frank Scholle^m^, Thiago Moreno L. Souza^d,e^, Nathaniel John Moorman^b,h,n^, Ralph S. Baric^b,c,n^, Peter Madrid^o^ and Sean Ekins^a#^

**Affiliations:** ^a^Collaborations Pharmaceuticals, Inc., 840 Main Campus Drive, Lab 3510, Raleigh, NC 27606, USA.

^b^Department of Microbiology and Immunology, University of North Carolina School of Medicine, Chapel Hill NC 27599, USA.

^c^Department of Epidemiology, University of North Carolina School of Medicine, Chapel Hill NC 27599, USA.

^d^Laboratório de Imunofarmacologia, Instituto Oswaldo Cruz (IOC), Fundação Oswaldo Cruz (Fiocruz), Rio de Janeiro, RJ, Brazil

^e^Centro De Desenvolvimento Tecnológico Em Saúde (CDTS), Fiocruz, Rio de Janeiro, Brasil

^f^Laboratory of Tropical Diseases – Prof. Dr. Luiz Jacinto da Silva, Department of Genetics, Evolution, Microbiology and Immunology, University of Campinas-UNICAMP, Campinas, SP, Brazil.

^g^Department of Microbiology and Immunology, University of Maryland School of Medicine, Baltimore, Maryland, USA

^h^Center for Integrative Chemical Biology and Drug Discovery, Chemical Biology and Medicinal Chemistry, Eshelman School of Pharmacy, University of North Carolina, Chapel Hill, North Carolina 27599, USA.

^i^UNC Lineberger Comprehensive Cancer Center, Chapel Hill, North Carolina 27599, USA.

^j^Institute for Antiviral Research, Utah State University, Logan, UT, USA.

^k^Department of Animal, Dairy and Veterinary Sciences, Utah State University, Logan, UT, USA.

^l^LabMol - Laboratory of Molecular Modeling and Drug Design, Faculdade de Farmácia, Universidade Federal de Goiás, Goiânia, GO, 74605-170, Brazil.

^m^Department of Biological Sciences, North Carolina State University, Raleigh, NC, USA.

^n^Rapidly Emerging Antiviral Drug Discovery Initiative, University of North Carolina at Chapel Hill, Chapel Hill, NC, USA.

^o^SRI International, 333 Ravenswood Avenue, Menlo Park, CA 94025, USA.

^#^ To whom correspondence should be addressed: Ana C. Puhl, E-mail address:; Sean Ekins, E-mail address:, Phone: +1 215-687-1320

**Figure S1.** **Dose response data from Vero E6 cells.** % inhibition is shown in blue and % cytotoxicity is shown in red. **A)** Pyronaridine**, B)** Tilorone and **C)** Quinacrine.

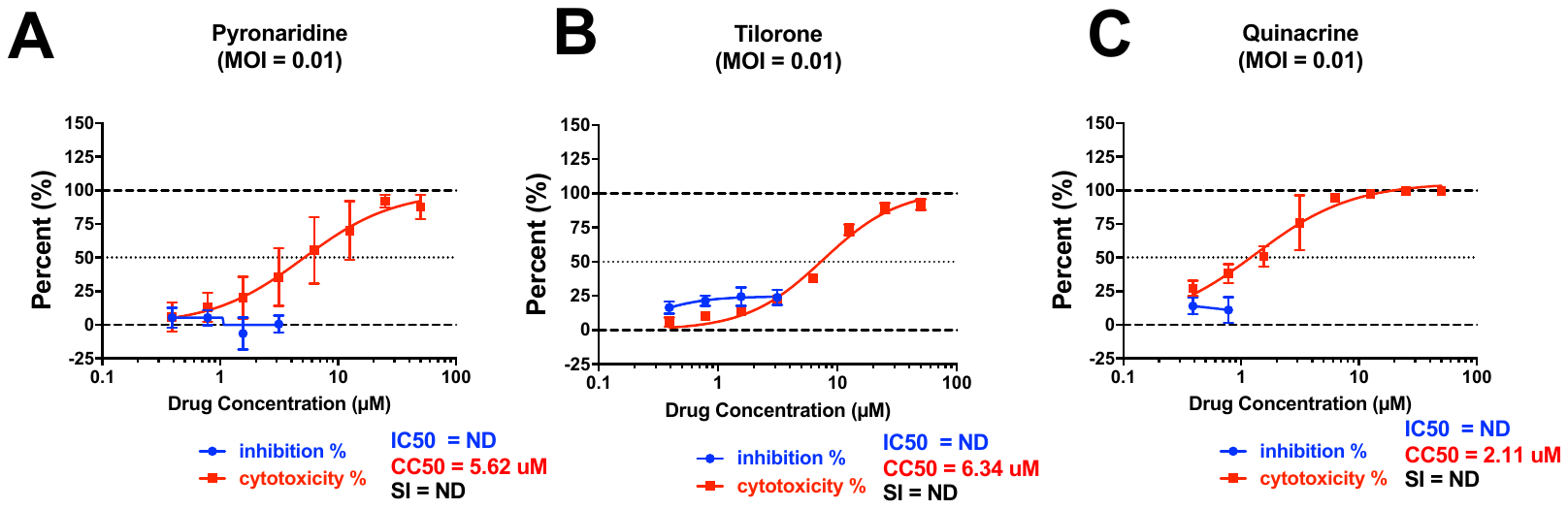

**Figure S2:** ***In vitro* antiviral SARS-CoV-2 testing in Calu-3 cells.** Calu-3 (ATCC, HTB-55) cells were pretreated with test compounds for 2 hours prior to continuous infection with SARS-CoV-2 (isolate USA WA1/2020) at a MOI=0.5. Forty-eight hours post-infection, cells were fixed, immunostained, and imaged by automated microscopy for infection (dsRNA+ cells/total cell number) and cell number. Sample well data was normalized to aggregated DMSO control wells and plotted versus drug concentration to determine the IC_50_ (infection: blue) and CC_50_ (toxicity: green). Percentage of Control (POC)=(sample well measurement /aggregated DMSO avg)*100 for n=3 replicates. A. Pyronaridine, B. Quinacrine, C. Tilorone, D. Remdesivir.

**Figure S3. Yield-reduction assays in different cell lines.** A. Human hepatoma lineage (HUH-7) were infected at MOI of 0.1 for 1 h at 37 °C and treated with different concentrations of tilorone. Lysis of cell monolayer was performed 48 h post infection and cell-associated viral RNA was quantified by real time RT-PCR. B. Tilorone impairs SARS-CoV-2 replication in human primary monocytes. Human primary monocytes were infected at the indicated MOI of 0.01 and treated with 10 µM of tilorone. After 24h, cell-associated viral RNA levels were measured by real time RT-PCR. C and D. Lung epithelial cell line (CALU-3) at MOI of 0.1 for 1 h at 37 °C and treated with different concentrations of tilorone. Lysis of cell monolayer was performed 48 h post infection and virus was titrated by plaque-forming units (PFU) assays and reported as % inhibition (C) and PFU/mL (D)**.**

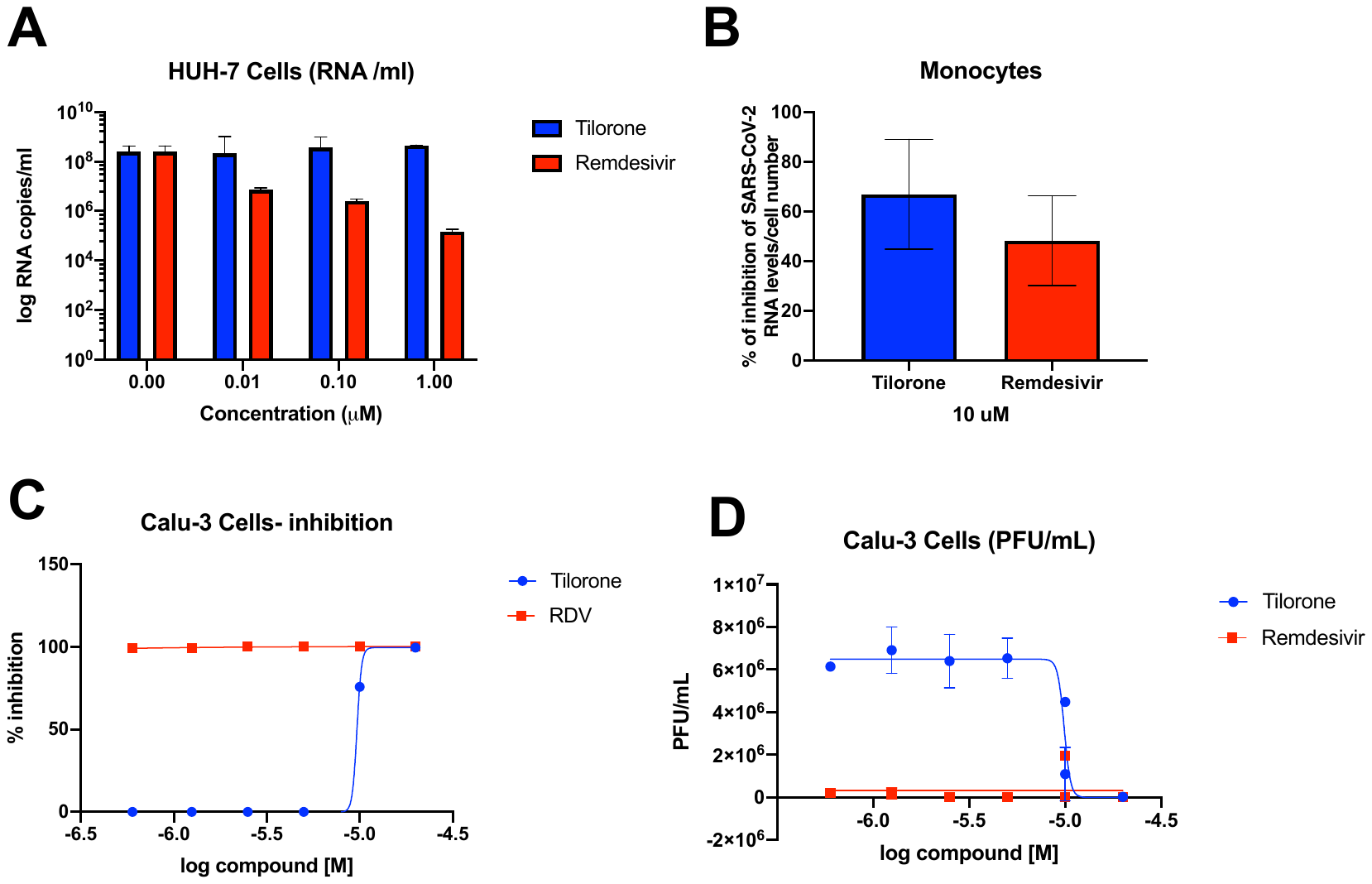

**Figure S4. Cytotoxicity assays in human monocytes and Vero CCL81 cells**. Human monocytes and/ or Vero CCL81 cells were incubated with different concentrations of the compounds A. tilorone, B. quinacrine and C. pyronaridine, for 24h and 72h respectively, for further cellular metabolic evaluation by measuring the activity of a mitochondrial enzyme succinate dehydrogenase using MTT test. CC50 values obtained for monocytes or Vero cells are described in blue and red respectively. One independent experiment was performed in duplicate.

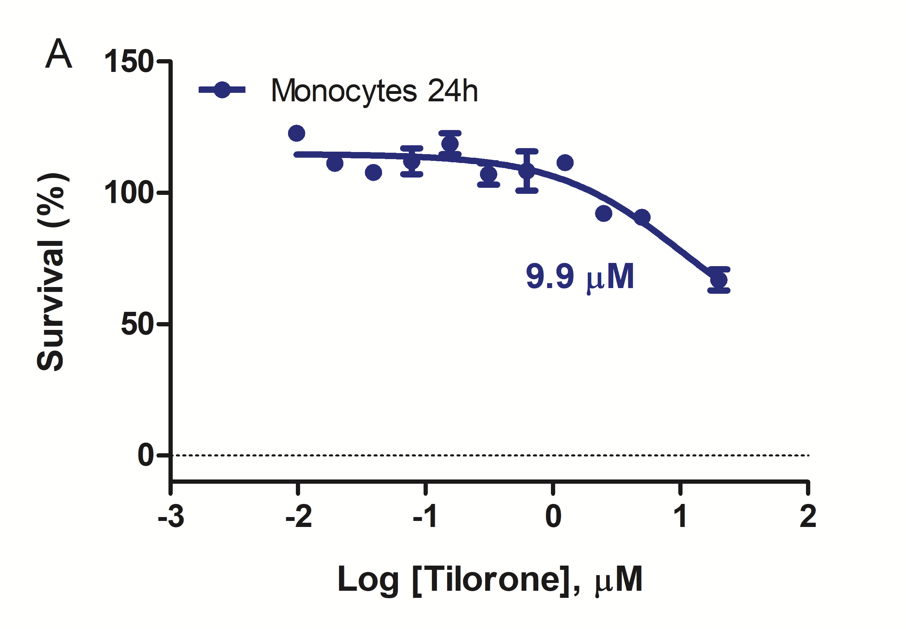

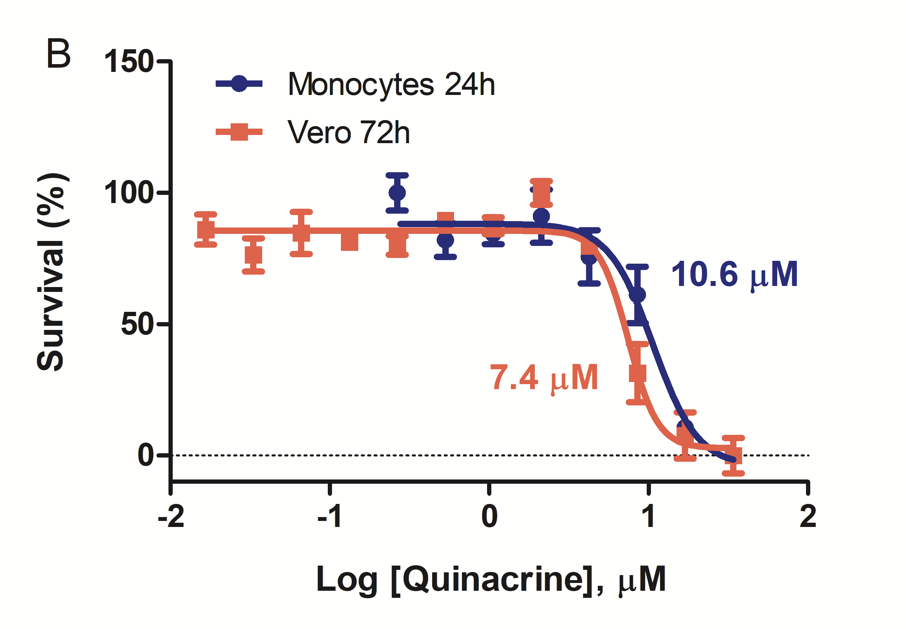

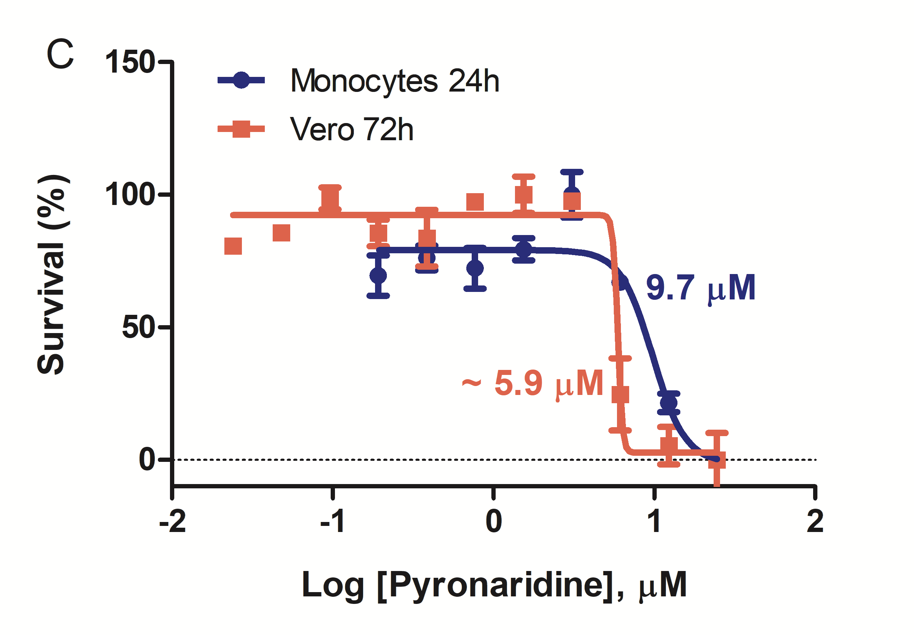

**Figure S5. Additional coronavirus testing.** A. HCoV229E antiviral assay and B. cytotoxicity in Huh-7 cell line.

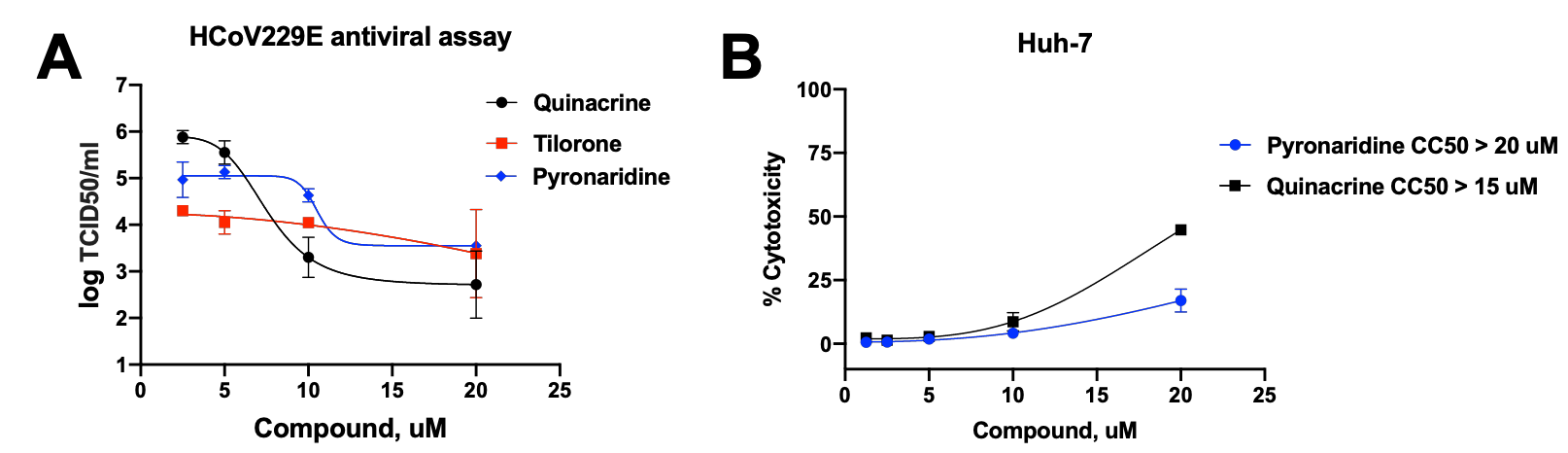

**Figure S6. Pseudovirus testing**. Neutralization activity for A. tilorone and B. pyronaridine. None of the concentrations tested reached 50% neutralization.

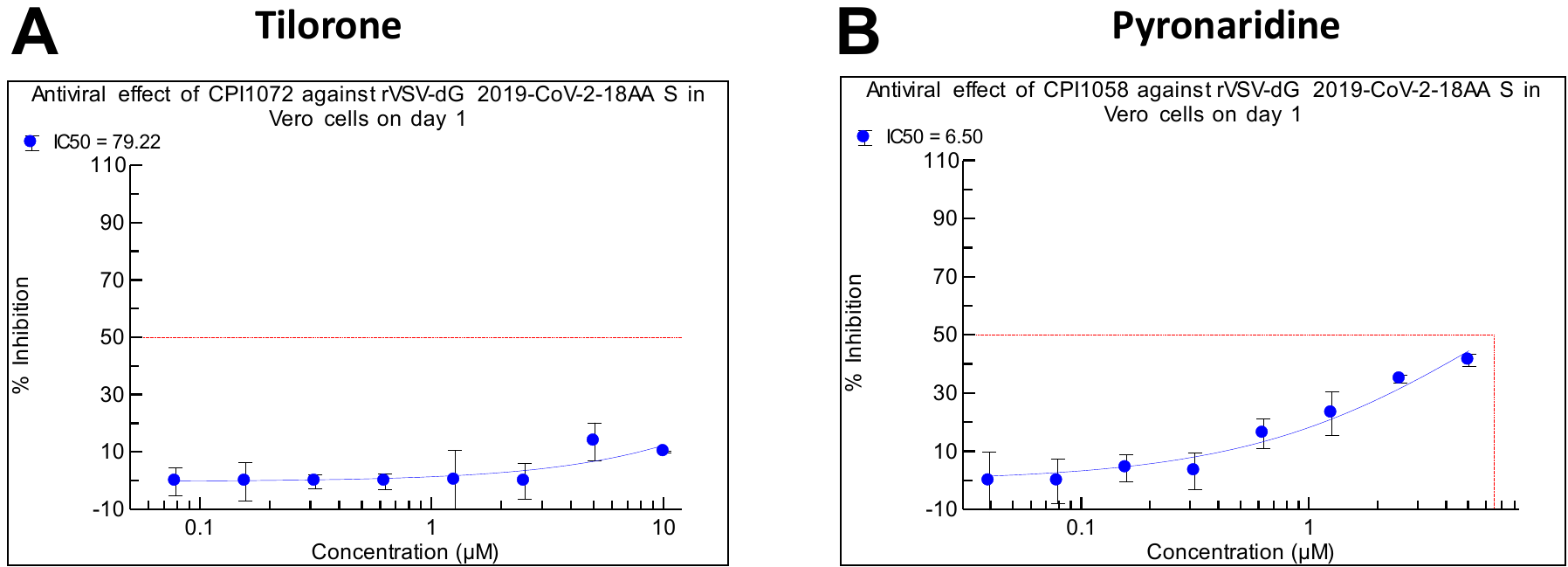

**Table S1**. Summary of all cell results infected with SARS-CoV-2, MHV and HCoV229E, X = inactive, Y = active, blank = not tested.

|  | Vero E6 | Vero 76 | Caco-2 | Calu-3 | A549-ACE2 | MHV | Huh-7 | Monocytes | HCoV229E | Pseudovirus |
| --- | --- | --- | --- | --- | --- | --- | --- | --- | --- | --- |
| Pyronaridine | X | X | Y | X | Y | Y |  |  | Y | weak |
| Tilorone | X | Y | Y | Y | Y | Y | X | Y | X | weak |
| Quinacrine | X | X | Y | X | Y | Y |  |  | Y |  |

**Table S2.** Summary of mouse pharmacokinetics and *in vitro* average Ebola IC_50_ (3 strains(Lane et al., 2020b)) *tilorone was dosed *in vivo* against Ebola at 25 mg/kg. Tilorone and pyronaridine data is from male mice only. Pharmacokinetics data from recent publications (Ekins et al., 2018b; Lane et al., 2019a; Lane et al., 2019b).

| **Drug** | **Dose (mg/kg)** | **C_max_ (ng/ml)** | **C_max_ (μM)** | **Ave IC_50_ (μM)** | **CC_50_ (μM)** | **Is the C_max_ above IC_50_** |
| --- | --- | --- | --- | --- | --- | --- |
| Quinacrine | 25 | 575* | 1.44 | 1.31 | 8.62 | Yes |
| Tilorone | 10 | 135 | 0.33 | 1.28 | 18.04 | No * |
| Pyronaridine | 75 | 982 | 1.9 | 1.04 | 8.62 | Yes |

*previously unpublished data

**Table S3.** Summary of human pharmacokinetics and *in vitro* average Ebola IC_50_ (3 strains (Lane et al., 2020b)). Pyronaridine pharmacokinetics from a single oral dose (Blessborn et al., 2017). Quinacrine pharmacokinetics from an intrapleural dose (Bjorkman et al., 1989).

| **Drug** | **Dose (mg)** | **C_max_ (ng/ml)** | **C_max_ (μM)** | **Ave IC_50_ (μM)** | **CC_50_ (μM)** | **Is the C_max_ above IC_50_** |
| --- | --- | --- | --- | --- | --- | --- |
| Quinacrine | 600 | 970 | 2.43 | 1.31 | 8.62 | Yes |
| Tilorone | 125 | unknown | Unknown | 1.28 | 18.04 | Unknown |
| Pyronaridine | 400 | 495.8 | 1.01 | 1.04 | 8.62 | Yes |

**Table S4.** *In vitro* and *In vivo* data for pyronaridine, quinacrine and tilorone (Ekins et al., 2018b; Lane et al., 2019a; Lane et al., 2019b; Lane et al., 2020b).

| **ADME property** | **Pyronaridine** | **Quinacrine** | **Tilorone** |
| --- | --- | --- | --- |
| **Solubility** | 168 µM at pH 7.4 | 80.5 µM at pH 7.4 | 465 µM at pH 7.4 |
| **CYP inhibition** | 1A2, 2C9, 2C19 (>50 µM), 3A4 (42.9 µM), 2D6 (2.23 µM) | 1A2 (6.66 µM), 2C9 (>50 µM), 2C19 (8.66 µM), 3A4 (3.96 µM), 2D6 (0.013 µM) | CYPs 1A2, 2C9, 2C19, 3A4, 2D6 (>50 µM) |
| **Mouse liver**  **microsomes** | t_1/2_ = >186 min,  CL_int_ = <7.4 µL/min/mg protein | t_1/2_ = 12.6 min,  Cl_int_= 110.0µL/min/mg protein | t_1/2_ = 102.7 min  CL_int_ = 13.5 µL/min/mg protein |
| **Guinea Pig liver microsomes** | t_1/2_ = 66.1 min,  CL_int_ = 21.0 µL /min/mg protein | t_1/2_ = 17.1min,  CL_int_ = 81.3µL /min/mg protein | t_1/2_ = 12.2 min, CL_int_ = 113.7 µL /min/mg protein |
| **Non-Human primate liver microsomes** | t_1/2_ = 89.7 min,  CL_int_ = 15.5 µL /min/mg protein | t_1/2_ = 10.1 min,  CL_int_ = 137.4 µL /min/mg protein | t_1/2_ = 94.3 min, CL_int_ = 14.7 µL /min/mg protein |
| **Human liver microsomes** | T_1/2_ = 122.2 min,  CL_int_ = 11.4 µL /min/mg protein | T_1/2_ = 27.5 min,  CL_int_ = 50.5 µL /min/mg protein | t_1/2_ 127.1 min, CL_int_ 10.9 µL/min/mg protein |
| **Mouse plasma protein binding** | 96.5% | 91.8% | 61.4 % |
| **Human plasma protein binding** | 95.1% | 89.9% | 52 % |
| **Caco-2** | Papp A-B = 6.46; B-A = 4.8 (x10^-6^ cm/s) Efflux ratio = 0.74 | P_app_ A-B = 30.6xE-6 cm/s  B-A = 19.0xE-6 cm/s  Efflux ratio = 0.92 | Papp A-B =20.4; B-A = 8.87(x10-6cm/s) Efflux ratio= 0.435 |
| **CYP3A4 Induction** | 1.5x at 10 µM | Not tested | Not tested |
| **Maximum Tolerated dose in mice (Balb/c)** | 100mg/kg | 50mg/kg* | 100 mg/kg |

*previously unpublished data
